# Structural Basis for eIF2B Inhibition in Integrated Stress Response

**DOI:** 10.1101/503540

**Authors:** Kazuhiro Kashiwagi, Takeshi Yokoyama, Madoka Nishimoto, Mari Takahashi, Ayako Sakamoto, Mayumi Yonemochi, Mikako Shirouzu, Takuhiro Ito

## Abstract

A core event in the integrated stress response, an adaptive pathway common to all eukaryotic cells in response to various stress stimuli, is the phosphorylation of eukaryotic translation initiation factor 2 (eIF2). Normally, unphosphorylated eIF2 transfers methionylated initiator tRNA to the ribosome in a GTP-dependent manner. In contrast, phosphorylated eIF2 inhibits its specific guanine nucleotide exchange factor eIF2B, which leads to a deficiency of active eIF2 and resultant global translation repression. To unveil the mechanism by which the eIF2 phosphorylation status regulates the eIF2B nucleotide exchange activity, we determined cryo-electron microscopic and crystallographic structures of eIF2B in complex with unphosphorylated or phosphorylated eIF2. Intriguingly, the unphosphorylated and phosphorylated forms of eIF2 bind to eIF2B in completely different manners: the nucleotide exchange-active “productive” and nucleotide exchange-inactive “nonproductive” modes, respectively. The nonproductive-mode phosphorylated eIF2, extending from one of the two eIF2B “central cavities”, not only prevents nucleotide exchange on itself, but also sterically prevents unphosphorylated eIF2 from productively binding on the other central cavity of eIF2B, which explains how phosphorylated eIF2 inhibits eIF2B.

**One Sentence Summary:** A drastic change in the binding mode of eIF2 to eIF2B induces translational control in stress.

---

Eukaryotic cells regulate protein synthesis in response to environmental stress. Various stress signals commonly induce the phosphorylation of eukaryotic translation initiation factor 2 (eIF2) in the stress-adaptive process known as the integrated stress response (ISR), which is conserved from yeast to human (1). eIF2 is a GTP-dependent carrier of methionylated initiator tRNA (Met-tRNA_i_) to the ribosomes, and activated by the eIF2-specific guanine nucleotide exchange factor (GEF) eIF2B (2). In mammals, four eIF2 kinases, GCN2, HRI, PERK, and PKR, respond to various types of stressors and phosphorylate eIF2 (3). The phosphorylation converts eIF2 from the substrate to an inhibitor of eIF2B, and thereby reduces the cellular level of active GTP-bound eIF2. The shortage of active eIF2 restricts the supply of Met-tRNA_i_ to the ribosomes, leading to global attenuation of translation initiation. While the eIF2 phosphorylation leads to global translation repression, it allows the selective translation of a specific subset of mRNAs required for cell survival and recovery, through alternative mechanisms such as reinitiation (4, 5). The ISR program is reportedly blocked by the small molecule ISRIB (ISR inhibitor), which enhances the nucleotide exchange activity of eIF2B (6, 7). This molecule has attracted a lot of interest because of its potential for the treatments of vanishing white matter disease, which is caused by mutations in the eIF2B subunits (8, 9), neurodegeneration (10), and traumatic brain injury (11). Therefore, ISR is induced or blocked depending upon the catalytic activity of eIF2B against eIF2.

eIF2 is a heterotrimeric protein consisting of α-, β-, and γ-subunits. eIF2 kinases phosphorylate the Ser51 residue of the α-subunit of eIF2 (eIF2α), and the γ-subunit holds GTP/GDP (3, 12). eIF2B is a decameric protein composed of two sets of five different subunits, α to ε (13). The X-ray crystallographic analysis of the overall structure of eIF2B revealed the central hexamer, composed of two sets of the α-, β-, and δ-subunits, which is sandwiched by two γε heterodimers (14). The γε heterodimer is mainly involved in the nucleotide exchange activity (15). The minimum catalytic core is the HEAT domain at the C-terminus of eIF2Bε, but the other part of eIF2B is required for efficient nucleotide exchange (15–17). The α2β2δ2 central hexamer is mainly involved in the recognition of phosphorylated eIF2α, and the “central cavity”, constructed by one set of the α-, β-, and δ-subunits and located at each pole of the central hexamer, performs this role (15, 18). Recent cryo-electron microscopic (cryo-EM) studies identified the binding site of ISRIB, and revealed that it bridges two δ-subunits and promotes the assembly of eIF2B (19, 20). Some mechanisms have been discussed, based on the eIF2B structure (14, 21), but the complex structure between eIF2 and eIF2B has not been solved. Therefore, the means by which eIF2B catalyzes nucleotide exchange on eIF2 and eIF2B is inhibited by the phosphorylation of eIF2 have remained enigmatic. To investigate the eIF2B-binding modes of unphosphorylated eIF2 and phosphorylated eIF2 [eΙF2(αP)], we performed a cryo-EM analysis.

## Cryo-EM structures of eIF2•eIF2B complexes

The complexes of eIF2•eIF2B or eIF2(αP)•eIF2B were prepared by mixing the purified proteins together with detergent, and analyzed by cryo-EM (figs. S1 and S2). The resultant structures of the eIF2•eIF2B and eIF2(αP)•eIF2B complexes revealed that the interaction mode between eIF2 and eIF2B is drastically changed by the phosphorylation of eIF2: the unphosphorylated and phosphorylated eIF2 extend in completely different ways from the eIF2B central cavity (Fig. 1 and fig. S3).

The structure of the unphosphorylated eIF2•eIF2B complex was determined at an overall resolution of 4.0 Å (Fig. 1A, figs. S1, S3A, and table S1). One molecule of eIF2 bound to the eIF2B decamer was clearly resolved in the cryo-EM map, and only a trace of density was observed on the opposite side of eIF2B. Even though the class containing two molecules of eIF2 was absent in this reconstruction, the size exclusion chromatography analysis of the eIF2•eIF2B mixture and the low-resolution map from the sample prepared without detergent revealed that two molecules of eIF2 can bind simultaneously to eIF2B (fig. S1, F and G). Therefore, the absence of one eIF2 molecule seems to be simply due to the scarcity of particles of the complex containing two molecules of eIF2, rather than a mechanism blocking such complex formation.

**Fig. 1.**
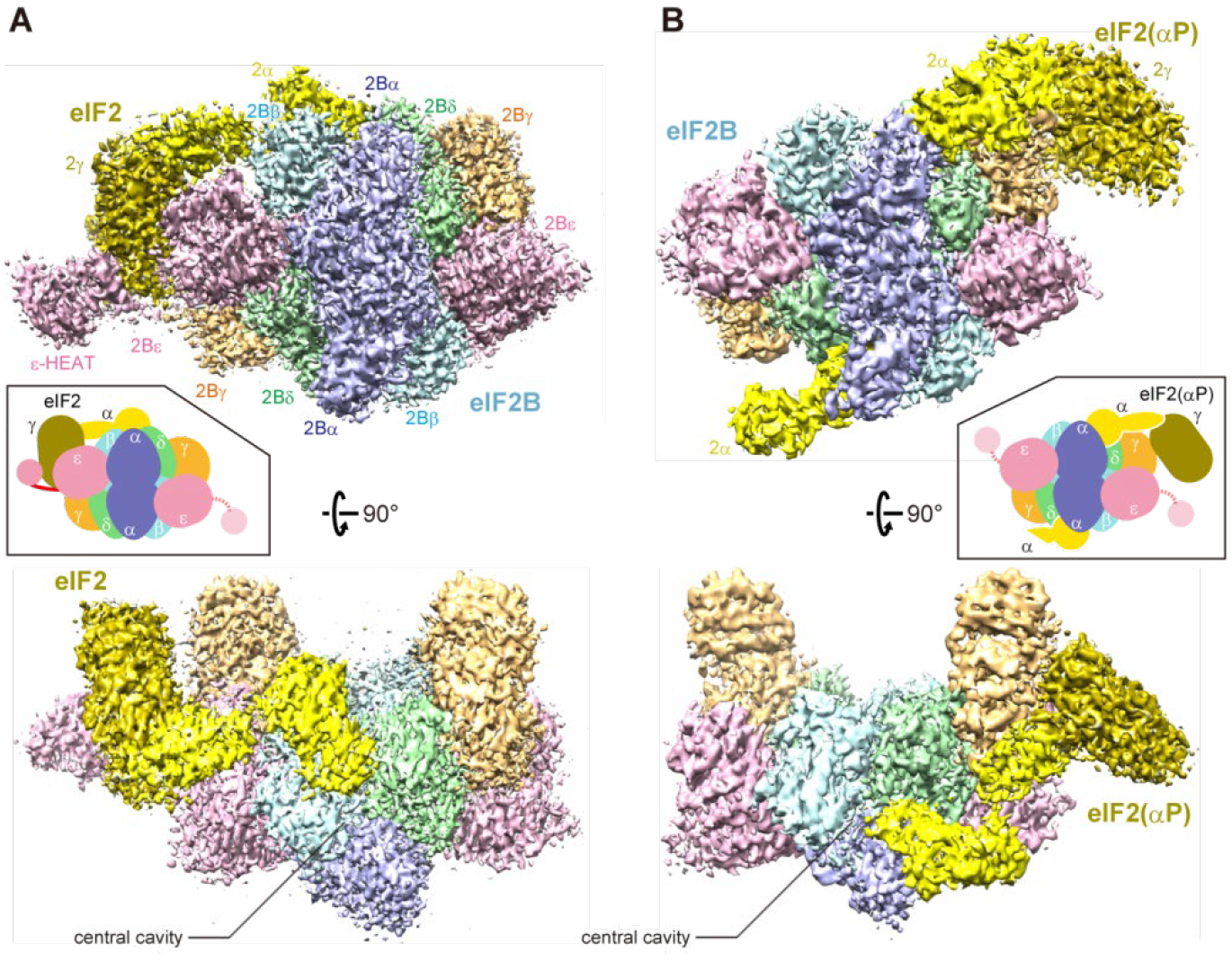
Cryo-EM maps of the human eIF2•eIF2B complexes. Two different views of the complexes of unphosphorylated eIF2•eIF2B (A) and phosphorylated eIF2(αP)•eIF2B (B). Subunit colors: eIF2α, yellow; eIF2γ, olive; eIF2Bα, purple; eIF2Bβ, cyan; eIF2Bγ, orange; eIF2Bδ, green; eIF2Bε, pink.

Similarly, the structure of the eIF2(αP)•eIF2B complex was determined at an overall resolution of 4.5 Å (Fig. 1B, figs. S2, S3B, and table S1). The α- and γ-subunits of eIF2(αP) were resolved on one side of eIF2B, and only the N-terminal domain (NTD) of eIF2α was resolved on the opposite side. As mentioned above, the binding region and the orientation of eIF2(αP) are completely different from those of the unphosphorylated eIF2.

In both structures, the cryo-EM density of eIF2β, which is supposed to be attached to eIF2γ, was not clearly observed because of its high mobility or loose association with eIF2γ, as suggested in the previous study (22). Importantly, only one of the two interaction modes was observed in the respective reconstructions. In both complexes, the accommodation of eIF2 did not cause large conformational changes of eIF2B.

## Catalysis of nucleotide exchange on eIF2

In the unphosphorylated eIF2•eIF2B complex, eIF2γ is sandwiched between the HEAT domain and the main body of eIF2Bε (Figs. 1A and 2A). This is the first structure in which the HEAT domain is observed together with the other parts of eIF2B. We regarded this structure as the “productive” complex, in which eIF2B catalyzes nucleotide exchange on eIF2. This structure is expected to represent the state after the dissociation of GDP from eIF2, since this sample was prepared without nucleotides. In this structure, the HEAT domain binds across the G domain and domain III of eIF2γ, and the NF motif, with the conserved consecutive Asn263–Phe264 residues of eIF2Bε (17), penetrates into eIF2γ from the main body side. Considering that the amino acid substitutions of these Asn and Phe residues reportedly reduce the catalytic activity to the level of the HEAT domain alone, these residues seem to fix eIF2γ in the favorable orientation for catalysis.

The N-terminal two helices of the HEAT domain interact with eIF2γ (Fig. 2A). These two helices in the HEAT domain contain most of the catalytically important residues identified by genetic and biochemical studies in *Saccharomyces cerevisiae* (23), and the most critical residue, Glu577 (Glu569 in *S. cerevisiae*) (24), directly interacts with eIF2γ. However, the interface for the HEAT domain in eIF2γ is separate from the nucleotide binding pocket, and this domain does not interact directly with the switch regions (Fig. 2A and fig. S4A). Therefore, the HEAT domain seems to catalyze the nucleotide exchange by inducing the structural rearrangement of eIF2γ. As compared with the structure of GDP-bound aIF2γ (25), the archaeal homolog of eIF2γ, the relative positions of domains II and III are slightly displaced by the interaction with the HEAT domain (fig. S4A). In the G domain, the most noticeable change is the conformation of the switch 1 region, which recognizes the phosphate moiety of the nucleotide (Fig. 2B and fig. S4B). The switch 1 region adopts a more open conformation in this productive complex, as compared with that in aIF2γ-GDP, which seems to make the primary contribution to the GDP dissociation. The opened switch 1 region extends toward the main body of eIF2Bε, close to some residues including Asn263 in the NF motif, suggesting an additional role of the NF-motif. Taken together, the cooperative action of the HEAT domain and the main body of eIF2Bε keeps the switch 1 region open, and induces the dissociation of GDP.

**Fig. 2.**
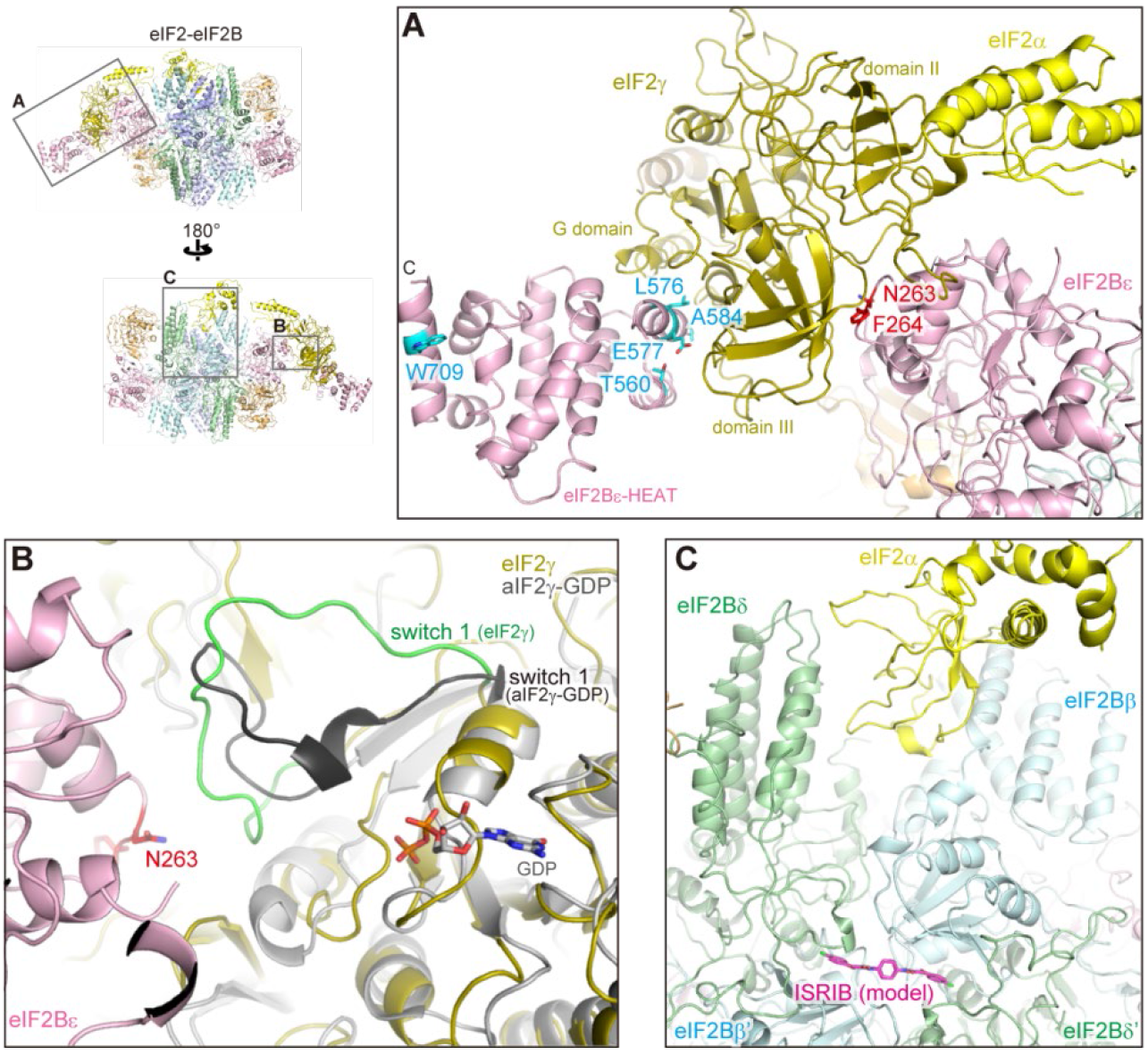
Interactions between unphosphorylated eIF2 and eIF2B. **(A)** The interaction between eIF2γ and eIF2Bε. The NF motif residues in eIF2Bε are colored red and the key residues for the catalytic function of eIF2B in the HEAT domain (23) are colored cyan. **(B)** Close-up view of the switch 1 region of eIF2γ (green). The GDP-bound aIF2γ structure from the aIF2-GDP structure (PDB ID: 2QMU) (25) is shown in grey and aligned to eIF2γ. The 3 10 helix in the switch 1 region (black) unfolds and extends toward the main body of eIF2Bε. **(C)** The interaction between the eIF2α and eIF2B subunits. The position of the ISRIB molecule in the human eIF2B-ISRIB complex structure (PDB ID: 6CAJ) (19) is shown in magenta.

In this productive binding mode, the NTD of eIF2α is recognized by the β- and δ-subunits of the eIF2B central cavity (Figs. 1A and 2C). The absence of an interaction between eIF2α and eIF2Bα was unexpected, because the amino acid substitution in eΓF2Bα abolished the eΓF2B•eIF2α interaction in our previous ITC experiment (14). Intriguingly, this interface for eIF2α is located just above the binding pocket for ISRIB (Fig. 2C) (19, 20). ISRIB bridges the two δ-subunits and promotes the assembly of eIF2B, from two βγδε tetramers and one α•α dimer into the decamer (19). Our structure revealed that the promotion of the assembly also stabilizes the productive interface for eIF2α. Therefore, ISRIB supports the stable interaction between eIF2 and eIF2B, enhancing the catalytic activity of eIF2B.

After GDP dissociates from eIF2γ, it is generally considered that GTP and Met-tRNA_i_ binding enables the detachment of eIF2 from eIF2B, stimulated by another initiation factor, eIF5 (26, 27). The structural alignment of eIF2γ in the productive complex and aIF2γ in the aIF2•GTP•Met-tRNA_i_ ternary complex (aIF2-TC) structure (28) revealed that Met-tRNA_i_ can access eIF2 in the productive complex, without clashing with most of eIF2B (fig. S4C). However, the path of the CCA tail of Met-tRNA_i_ is blocked by the opened switch 1 region of eIF2γ in this structure (fig. S4D). GTP binding closes the switch 1 region of eIF2γ to facilitate the accommodation of Met-tRNA_i_. Following the correct accommodation of Met-tRNA_i_, the NTD of eIF2α moves from its interface on eIF2B toward Met-tRNAi and fits along its L-shape (fig. S4E). Thus, eIF2 seems to reduce the affinity for eIF2B and detaches from it.

## Recognition of eIF2 phosphorylation

In the eIF2(αP)•eIF2B complex, the binding mode of eIF2(αP) is different from that of the unphosphorylated eIF2. The NTD of the phosphorylated eIF2α (P-eIF2α) resides in between the α- and δ-subunits of the eIF2B central cavity, and eIF2γ docks onto eIF2Bγ (Fig. 1B and Fig. 3A). In this complex, eIF2γ does not interact with eIF2Bε, and the HEAT domain is not located at a fixed position. Therefore, this structure seems to represent the “nonproductive” complex, in which the nucleotide exchange reaction is restricted.

**Fig. 3.**
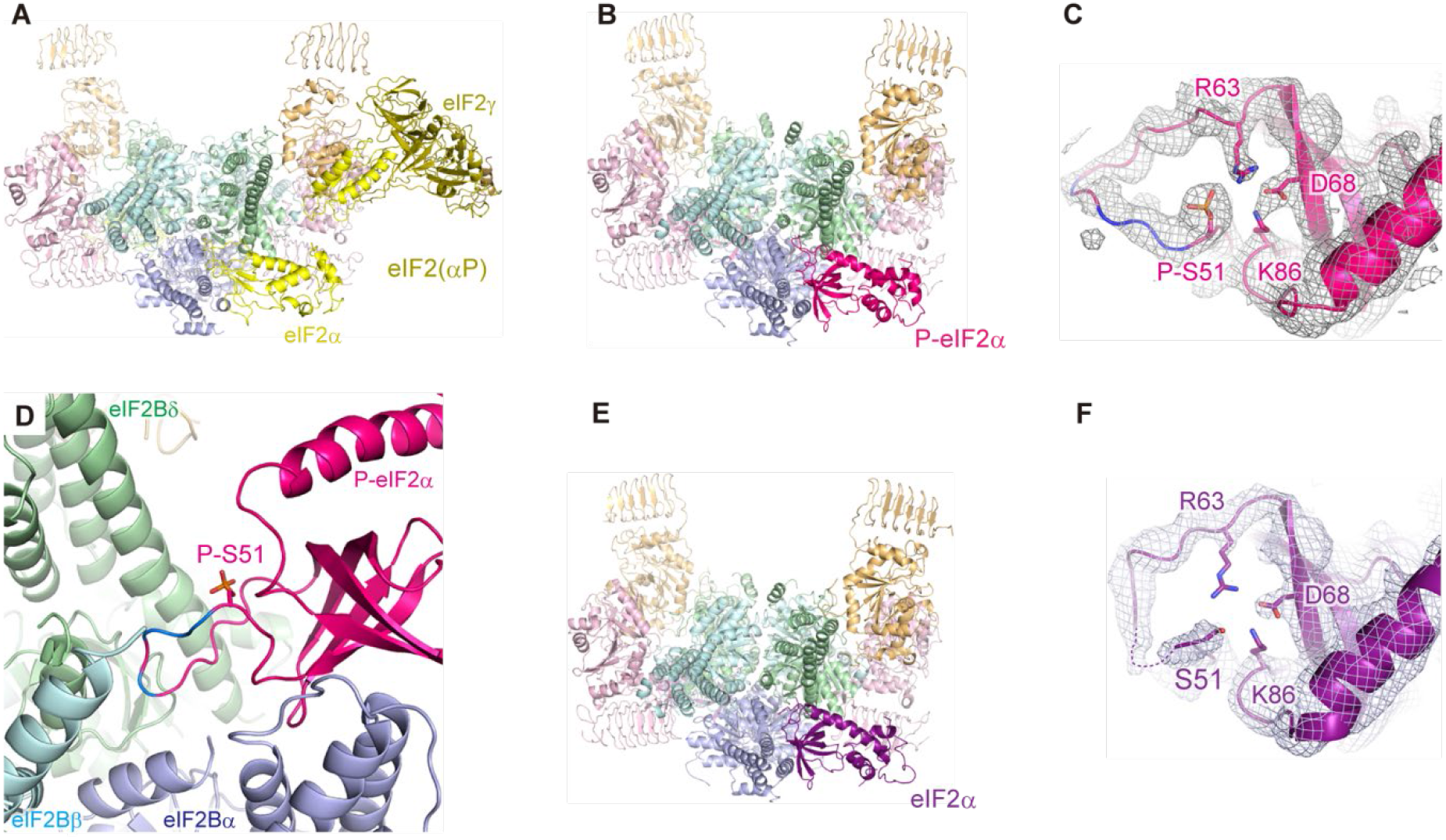
Recognition of phosphorylated eIF2α. (**A, B**, and **E**) The cryo-EM structure of the human eIF2(αP)•eIF2B complex (A), and the crystal structures of *S. pombe* eIF2B in complex with *S. cerevisiae* P-eIF2α (B) and eIF2B in complex with unphosphorylated eIF2α (E). (**C** and **F**) The electron density around the NTD of P-eIF2α (C) and eIF2α (F). The residues forming the electrostatic network together with P-Ser51 are shown in stick models. (**D**) Close-up view of the interface for P-eIF2α. The P-Ser51 residue is shown in a stick model and the arginine residues in the Ser51-flanking loop are colored blue.

P-eIF2α is mainly recognized by eIF2Bα, and a loop structure extends from the NTD of P-eIF2α toward the center of the eIF2B central cavity (fig. S5). This loop corresponds to the arginine rich region adjacent to the phosphorylated residue Ser51 (P-Ser51), but its resolution is insufficient to assign P-Ser51 and the flanking residues unambiguously.

To visualize the recognition mechanism of P-eIF2α in more detail, we determined the crystal structure of the complex of *S. cerevisiae* P-eIF2α and *Schizosaccharomyces pombe* eIF2B at 3.35-Å resolution (Fig. 3B and table S2). Even though only the NTD of P-eIF2α is resolved, it binds to eIF2B in a similar orientation to the above nonproductive complex, and the electron density for the P-Ser51 residue is clearly visible (Fig. 3C). As compared with the position of P-eIF2α-NTD in this crystal structure, that in the eIF2(αP)•eIF2B complex is slightly shifted toward eIF2α-CTD and eIF2γ (fig. S6). Therefore, P-eIF2α in the eIF2(αP)•eIF2B complex seems to be in a somewhat retracted position, presumably due to the association between eIF2γ and eIF2Bγ, whereas the P-eIF2α-NTD in the crystal structure is located where its phosphorylation status is examined. The P-Ser51 residue of P-eIF2α in the P-eIF2α•eIF2B complex is not directly recognized by eIF2B. Instead, this residue is involved in intramolecular electrostatic interactions with Arg63, Asp68, and Lys86 (Fig. 3C). The Ser51-flanking arginine residues protrude into the acidic part of the eIF2B central cavity, and interact with eIF2Bδ and the C-terminus of eIF2Bα (Fig. 3D).

A previous study has shown that some amino acid substitutions in eIF2α abrogate both eIF2B binding and phosphorylation by PKR, suggesting that eIF2B and PKR recognize overlapping surfaces on eIF2α (29). A comparison between our structure and the eIF2α-PKR complex structure (30) revealed that eIF2Bα and PKR recognize a similar region of eIF2α, and their manners of eIF2α recognition are also comparable, despite the lack of similarity in their amino acid sequences or overall tertiary structures (fig. S7, A to C). The angles and positions of the helices recognizing eIF2α (α3 of eIF2Bα and αG of PKR) are the same, and some residues form similar interactions with eIF2α: Lys79 of eIF2α interacts with Glu57 of eIF2Bα or Glu490 of PKR, and Asp83 of eIF2α caps the helix α3 of eIF2Bα or αG of PKR by interactions with the amide groups of their main chains (fig. S7, B and C).

We also obtained crystals of the complex of unphosphorylated eIF2α and eIF2B under similar crystallization conditions, and determined the structure at 3.5-Å resolution (Fig. 3E and table S2). This crystal structure revealed that unphosphorylated eIF2α binds to eIF2B in the same way as P-eIF2α. The only structural difference between them is limited to the neighbor of the Ser51-flanking loop. The loop region of unphosphorylated eIF2α is not well resolved and the protrusion of the loop is less prominent, as compared with that of P-eIF2α (Fig. 3F), in agreement with the fact that P-eIF2α interacts with eIF2B more tightly (31). The microscale thermophoresis measurement of the affinity between *S. cerevisiae* eIF2α and *S. pombe* eIF2B also confirmed that the phosphorylation of wild type eIF2α strengthened the affinity to eIF2B. The double alanine substitutions of Arg63 and Lys86, which participate in the intramolecular interaction network in P-eIF2α, abolished this phosphorylation-dependent affinity enhancement (fig. S7, D to F). These Ser51-interacting residues seem to work like a switch. They form a specific conformation through the electrostatic interaction network only when Ser51 is phosphorylated, and bind strongly to eIF2B by the deep insertion of the adjacent arginine-rich loop.

## Mechanism of eIF2B inhibition

The similar binding modes of eIF2α to eIF2B in the absence of eIF2β and -γ, regardless of its phosphorylation status, suggest that both the productive and nonproductive binding states are transiently allowed for the unphosphorylated eIF2, and the nonproductive binding is isomerized to more stable productive binding. In the case of eIF2(αP), such isomerization should be prevented. In the cryo-EM structure of the eIF2•eIF2B complex, Ser51 of eIF2α faces Glu139 of eIF2Bβ (Fig. 4A). Overlaying the P-eIF2α-NTD structure from the P-eIF2α•eIF2B complex revealed that the P-Ser51 residue comes closer to Glu139 of eIF2Bβ, and therefore, P-eIF2α is likely to be rejected from this interface due to electrostatic and steric repulsion (fig. S8A). This proposed rejection mechanism agrees with our previous cross-linking experiments between eIF2α and eIF2B (14). The cross-linked residues with unphosphorylated eIF2α are located around both the productive and nonproductive interfaces for eIF2α, and the cross-links around the productive interface were weakened by the phosphorylation of eIF2α (fig. S8B). In addition, the double alanine mutations at Glu164 and Leu165 in *S. cerevisiae* eIF2Bβ (Glu143 and Leu144 in human), which reside in the productive interface for eIF2α, partly mimic the eIF2α phosphorylation and enhance the nonproductive binding with unphosphorylated eIF2 (Fig. 4A) (32). These findings indicate that the abrogation of the interaction at the productive interface for eIF2α, by phosphorylation or mutations, converts the complex to the nonproductive binding mode, and inhibits nucleotide exchange.

**Fig. 4.**
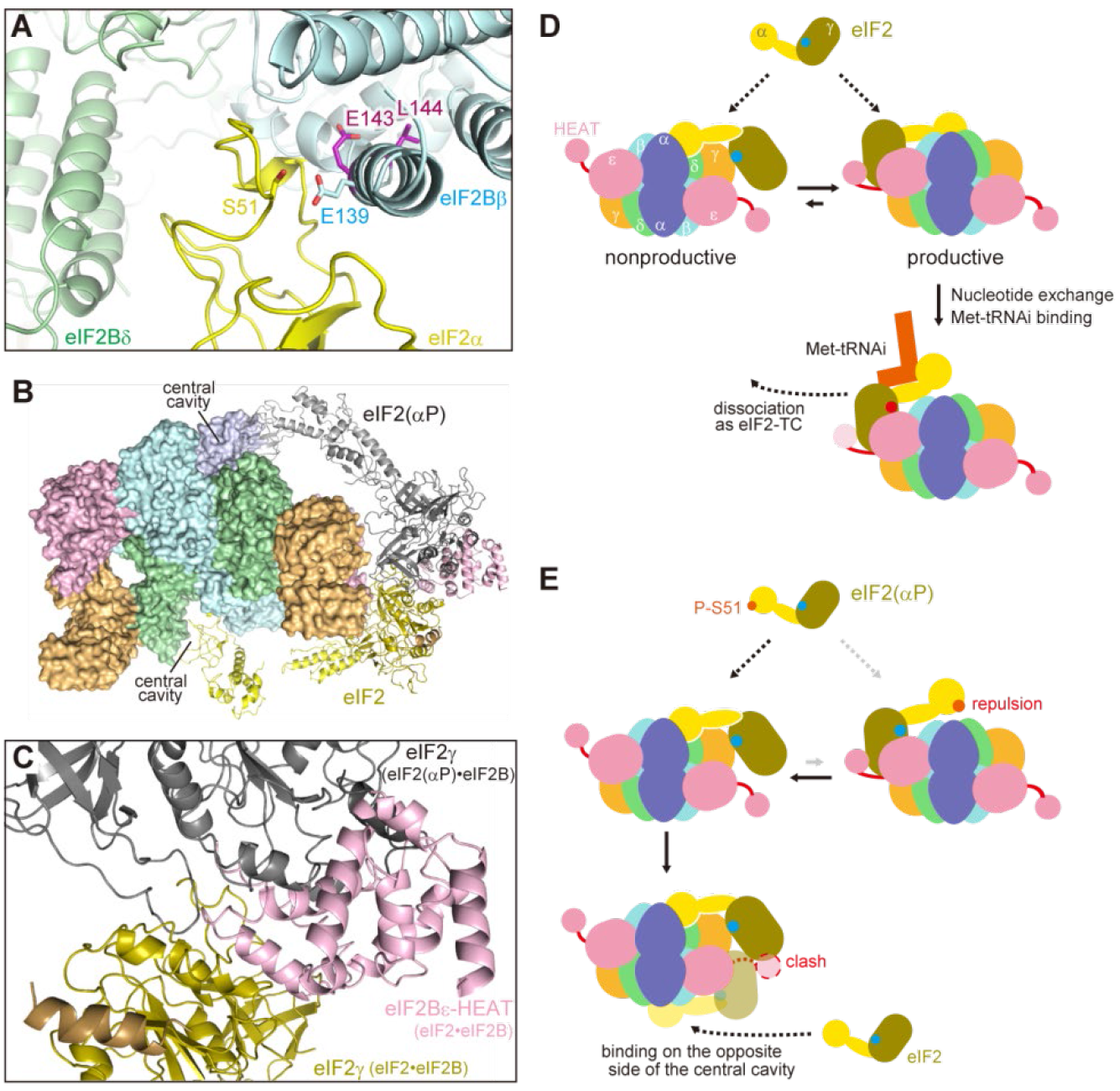
Mechanism of eIF2B inhibition by eIF2 phosphorylation. (**A**) Close-up view of the interface for eIF2α in the eIF2•eIF2B complex. Ser51 of eIF2α and Glu139 of eIF2Bβ are shown in stick models. The residues whose mutations partially mimic eIF2α phosphorylation in *S. cerevisiae* eIF2B are shown as purple sticks. (**B**) The structures of the eIF2•eIF2B and eIF2(αP)•eIF2B complexes are superimposed so that the two α subunits of eIF2 are bound on the opposite sides of the central cavity of eIF2B. eIF2(αP) is shown in grey. (**C**) Close-up view of the interface between eIF2γ and eIF2Bε-HEAT of the eIF2•eIF2B complex in the superimposed model. eIF2γ of the eIF2(αP)•eIF2B complex clashes with eIF2Bε-HEAT and slightly with eIF2γ of the eIF2•eIF2B complex. (**D** and **E**) Proposed mechanism of eIF2B inhibition by eIF2 phosphorylation. (D) The unphosphorylated eIF2 can bind to eIF2B in the productive or nonproductive mode and then shifts toward the more stable productive mode. A GDP molecule (cyan dot) bound to eIF2γ dissociates in this productive mode. GTP (red dot) binds and then Met-tRNAi binding induces the dissociation from eIF2B as eIF2-TC. (E) The mode of eIF2(αP) binding to eIF2B is limited to the nonproductive mode due to the repulsion at the productive interface for eIF2α, and nucleotide exchange is inhibited in this binding mode. The bound eIF2(αP) molecule also blocks the productive binding of eIF2 on the opposite side of eIF2B.

The binding of eIF2(αP) in the nonproductive mode not only prevents nucleotide exchange on itself but also inhibits the nucleotide exchange activity on the opposite side of the twofold symmetric eIF2B. In the nonproductive eIF2(αP)•eIF2B complex, eIF2γ docks on eIF2Bγ. If eIF2γ and the eIF2Bε HEAT domain were positioned as in the productive mode utilizing the opposite-side central cavity of eIF2B, then their positions would partially overlap with that of the near-side eIF2γ in the nonproductive complex (Fig. 4, B and C). Therefore, the binding of eIF2(αP) blocks the formation of the productive complex on the opposite side of eIF2B (Fig. 4, D and E).

In conclusion, our cryo-EM and crystal structures of the eIF2•eIF2B and eIF2(αP)•eIF2B complexes revealed that eIF2 has two distinct eIF2B-binding modes, and changes the favorable binding mode depending on the phosphorylation status of eIF2. The productive eIF2•eIF2B complex showed that the nucleotide exchange by eIF2B is performed by the concerted actions of multiple elements in the eIF2B decamer. The nonproductive eIF2(αP)•eIF2B complex indicated that the phosphorylated eIF2 inhibits the catalytic activity of eIF2B by remaining distant from the catalytic elements on eIF2B, and by blocking the formation of the productive complex with unphosphorylated eIF2 on the opposite side of eIF2B. This study provides a structural basis for the eIF2B inhibition by phosphorylated eIF2, the core mechanism of the integrated stress response.

## Acknowledgments

### Funding

This research was supported by Grants-in-Aid for Scientific Research on Innovative Areas “Nascent Chain Biology” (JP15H01548 and JP17H05677 to T.I.), a Grant-in-Aid for Scientific Research (B) (JP16H04756 to T.I.), and Grants-in-Aid for Early-Career Scientists (JP18K14644 to K.K.) from the Japan Society for the Promotion of Science (JSPS), Basis for Supporting Innovative Drug Discovery and Life Science Research (BINDS) (JP18am0101082) from the Japan Agency for Medical Research and Development (AMED), and the RIKEN Pioneering Project “Dynamic Structural Biology”. The data acquisitions of the crystal were performed under the approval of the Japan Synchrotron Radiation Research Institute (Proposals 2014A1347, 2014B1566, 2015B2050, 2016A2545, 2017A2851, and 2017B2851)

### Author contributions

K.K. and T.I. designed and performed experiments, and wrote the manuscript. T.Y. and M.S. performed the cryo-EM analysis. M.N., M.T., A.S., M.Y., and M.S. prepared the samples.

### Competing interests

The authors declare no competing financial interests.

### Data and materials availability

The structural coordinates of the eIF2•eIF2B, eIF2(αP)•eIF2B, P-eIF2α•eIF2B and eIF2α•eIF2B complexes have been deposited in the Protein Data Bank (PDB). The cryo-EM maps of the eIF2•eIF2B and eIF2(αP)•eIF2B complexes have been deposited in the Electron Microscopy Data Bank (EMDB). All other data can be obtained from the corresponding authors upon reasonable request.

