## Supplementary Materials for "Structural Basis for eIF2B Inhibition in Integrated Stress Response"

This PDF file includes:

Materials and Methods

Figs. S1 to S8

Tables S1 to S2

### Materials and Methods

No statistical methods were used to predetermine sample size. The experiments were not randomized. The investigators were not blinded to allocation during experiments and outcome assessment.

Purification of human eIF2

The α-, β-, and γ-subunits of human eIF2, and the eIF2-specific chaperone protein human Cdc123 (33) were co-expressed in FreeStyle 293-F cells, using the four pEBMulti-Neo plasmid vectors (Wako), and eIF2γ was expressed with C-terminal FLAG and $His_8$ tags. The cells were lysed in buffer A [20 mM MES-KOH buffer (pH 6.0), containing 150 mM KCl, 1 mM $MgCl_2$, 10%(v/v) glycerol, and 5 mM 2-mercaptoethanol] supplemented with 20 mM imidazole, 0.5 mM EDTA, 0.1%(v/v) Triton X-100 and protease inhibitors. After 30 min on ice and centrifugation, the supernatant was applied to a HisTrap (GE Healthcare) column equilibrated with buffer A supplemented with 20 mM imidazole, and eluted with a linear gradient of 20–500 mM imidazole. The fraction containing eIF2 was collected and applied to a HiTrap SP (GE Healthcare) column equilibrated with buffer A, and eluted with a linear gradient of 200–640 mM KCl. After three-fold dilution with buffer B [20 mM HEPES-KOH buffer (pH 7.5) containing 100 mM KCl, 0.1 mM $MgCl_2$, 10%(v/v) glycerol, and 1 mM DTT], the sample was applied to a HiTrap Heparin (GE Healthcare) column equilibrated with buffer B, and eluted with a linear gradient of 0.2–1 M KCl. The fraction containing eIF2 was further purified on a Superdex200 (GE Healthcare) column equilibrated with buffer B. eIF2 was phosphorylated by PKR, as described for *Komagataella pastoris* eIF2 (14).

Purification of human eIF2B

Human eIF2Bα and eIF2Bβγδε were purified separately.

The fragment encoding human eIF2Bα was cloned into pET-28c (Novagen), in which the thrombin cleavage site was replaced by the HRV 3C protease cleavage site. The T7 Express

Escherichia coli strain (New England Biolabs) was transformed with this plasmid and grown in LB medium at 37 °C. After the addition of 0.3 mM isopropyl-β-D-thiogalactopyranoside (IPTG) when the culture reached an absorbance, $A_{600\ nm}$, of 0.6, the cells were grown at 18 °C overnight. The cells were harvested and lysed by sonication in lysis buffer [20 mM HEPES-KOH buffer (pH 7.5), containing 150 mM KCl, 10%(v/v) glycerol, 1 mM DTT, and protease inhibitors]. After centrifugation, the supernatant was purified with cOmplete His-Tag Purification Resin (Roche), and passed through Q and SP Sepharose (GE Healthcare) resins. The HRV 3C protease was added to the sample and dialyzed against buffer C [20 mM HEPES-KOH buffer (pH 7.5), containing 150 mM KCl, 5%(v/v) glycerol, and 1 mM DTT] at 4 °C overnight. The sample was passed through Ni resin, and then purified on a Sephacryl S-300 column (GE Healthcare) in buffer C.

For the purification of eIF2Bβγδε, the fragments encoding eIF2Bβ and eIF2Bδ were cloned into pETDuet-1 (Novagen), and the fragments encoding eIF2Bγ and eIF2Bε were cloned into pCOLADuet-1 (Novagen). An HRV 3C protease-cleavable His6-tag was added to the N-terminus of eIF2B ε . The E. coli cells were transformed with these plasmids and grown as above. Cells were lysed by sonication in lysis buffer and centrifuged. The supernatant after centrifugation was purified with Ni Sepharose (GE Healthcare). The sample was applied to a HiTrap Heparin (GE Healthcare) column and eluted with a linear gradient of 0.15–1 M KCl. The fraction containing all four subunits was collected, and dialyzed against buffer C at 4 °C overnight with the HRV 3C protease. The sample supplemented with 10 mM imidazole was passed through the Ni resin, and then applied to a HiTrap Q (GE Healthcare) column and eluted with a linear gradient of 0.15–0.5 M KCl. After buffer exchange to buffer C using Amicon Ultra centrifugal filters (Millipore), eIF2Bβγδε were mixed with eIF2Bα at the molar ratio of 1:3 and applied to a Sephacryl S-300 (GE Healthcare) column equilibrated with buffer C.

Cryo-EM data acquisition and image processing

Human eIF2B, eIF2, and eIF2(αP) were respectively applied to a Superose 6 Increase (GE Healthcare) column equilibrated with 20 mM HEPES-KOH buffer (pH 7.5), containing 150 mM KCl, 5 mM $MgCl_2$, and 1 mM DTT, and the peak fractions were collected. eIF2B and eIF2 or eIF2(αP) were mixed at a molar ratio of 1:3. The samples were diluted to 100 nM for eIF2B and 300 nM for eIF2 or eIF2(αP), and supplemented with 0.06% digitonin just before application to glow-discharged Quantifoil R1.2/1.3 Holey Carbon Grids coated with carbon film. Using a Vitrobot Mark IV (FEI) at 4 °C and 100% humidity, 3 μl of samples were incubated on the grids for 30 s, blotted for 3 s, and plunged into liquid ethane. The datasets were collected with a Tecnai Arctica transmission electron microscope (FEI) operated at 200 kV, using a K2 Summit direct electron detector (Gatan) operated in the counting mode (1.47 Å/pixel). The data collection was automated with the SerialEM software (34), and the total numbers of collected images were 3,949 for the eIF2•eIF2B complex and 4,987 for the eIF2(αP)•eIF2B complex. The collected images were fractionated to 40 frames, with a total dose of ~50 $e^-/Å^2$ (table S1).

For the eIF2•eIF2B complex, the movie frames were aligned with MotionCor2 (35) and the CTF parameters were estimated with CTFFIND-4.1 (36) in RELION-3.0 (37). Manually picked particles were averaged in 2D, and the resulting classes were used as templates for automated particle picking with Gautomatch (http://www.mrc-lmb.cam.ac.uk/kzhang/). After 2D classification, 3D classification was performed using a low-pass filtered (40 Å) map calculated from the crystal structure of *S. pombe* eIF2B (PDB ID: 5B04) (14). The class containing eIF2 was further refined with 3D refinement, CTF refinement, and Bayesian polishing in RELION-3.0 (fig. S1C).

For the eIF2(αP)•eIF2B complex, image processing was performed as above, using RELION-2.1 (38). After 3D classification, the classes containing additional density were selected and subjected to 3D classification again. The classes containing eIF2α and eIF2 γ were selected and 3D refinement was performed. Then, the movie frames were re-processed with MotionCor2 and CTFFIND-4.1 in the RELION-3.0 package, and particles were extracted from the re-processed micrographs using the coordinates after the 3D refinement in RELION-2.1. Further refinement by 3D refinement, CTF refinement, and Bayesian polishing was performed using RELION-3.0 (fig. S2C).

The cryo-EM structure of human eIF2B in complex with ISRIB at 2.8-Å resolution (PDB ID: 6CAJ) (19), the poly-alanine model of the eIF2Bγ left-handed β-helix domain from the cryo-EM structure of human eIF2B in complex with ISRIB at 4.1-Å resolution (PDB ID: 6EZO) (20), and eIF2 from the cryo-EM structure of the human 48S pre-initiation complex (PDB ID: 6FEC) (39) were manually fitted to the density. The resulting models were refined using phenix.real_space_refine (40), and manually refined with Coot (41). The refinement statistics of these structures are shown in table S1.

Crystallization and structure determination

*S. pombe* eIF2B was purified as described (42). *S. cerevisiae* eIF2α was purified and phosphorylated by PKR, using the same protocol as described for *S. pombe* eIF2α (14). eIF2B and eIF2α were mixed at a molar ratio of 1:6, and concentrated to 10 μM for eIF2B and 60 μM for eIF2α. Crystals were grown at 20 °C by the sitting drop vapor diffusion method, by mixing with reservoir solution [150 mM sodium acetate, 75 mM sodium citrate (pH 5.7–6.4), 5–7% PEG 4000, and 5%(v/v) glycerol] at a 1:1 ratio. After a week, the crystals were cryoprotected by serial transfers into reservoir solutions supplemented with 5%(v/v), 10%, 15%, 20%, and 30% 1,3-propanediol, and flash-cooled in liquid nitrogen. Data collection was performed at BL41XU of SPring-8 (Hyogo, Japan). All datasets were collected at 100 K, at a wavelength of 1 Å.

The data sets were processed with XDS (43), and the initial phases were determined by molecular replacement with PHENIX (40), using the crystal structure of *S. pombe* eIF2B (PDB ID: 5B04) (14). Both of the resulting maps for the P-eIF2α•eIF2B and eIF2α•eIF2B complexes clearly revealed the electron densities for the two NTDs of eIF2α. The crystal structures of *S. cerevisiae* eIF2α-NTD (PDB ID: 1Q46) (44) were manually fitted in Coot. These structures were refined by iterative rounds of manual adjustment with Coot and refinement with PHENIX. The refinement statistics of these structures are shown in table S2. For the P-eIF2α•eIF2B complex, 88.5% of the residues in the model are in the favored region, 10.2% are in the allowed region, and 1.3% are in the disallowed region in the Ramachandran plot. For the eIF2α•eIF2B complex, 86.9% of the residues in the model are in the favored region, and 11.4% and 1.7% of the residues are in the allowed and disallowed regions, respectively.

Microscale thermophoresis

The R63A K86A substitutions of *S. cerevisiae* eIF2α were introduced using a PrimeSTAR Mutagenesis Basal Kit (Takara).

*S. pombe* eIF2B, and *S. cerevisiae* eIF2α and P-eIF2α were applied to gel filtration columns equilibrated with 20 mM HEPES-KOH buffer (pH 7.5), containing 150 mM KCl, 5 mM MgCl2, and 5%(v/v) glycerol. eIF2B was concentrated to 20 µM, and eIF2α or P-eIF2α was labelled with a fluorescent dye using a Monolith His-Tag Labeling Kit RED-tris-NTA (NanoTemper). The microscale thermophoresis analysis was performed with a Monolith NT.115 (NanoTemper) unit, according to the manufacturer's protocol. Each experiment was repeated three times for the calculation of dissociation constants.

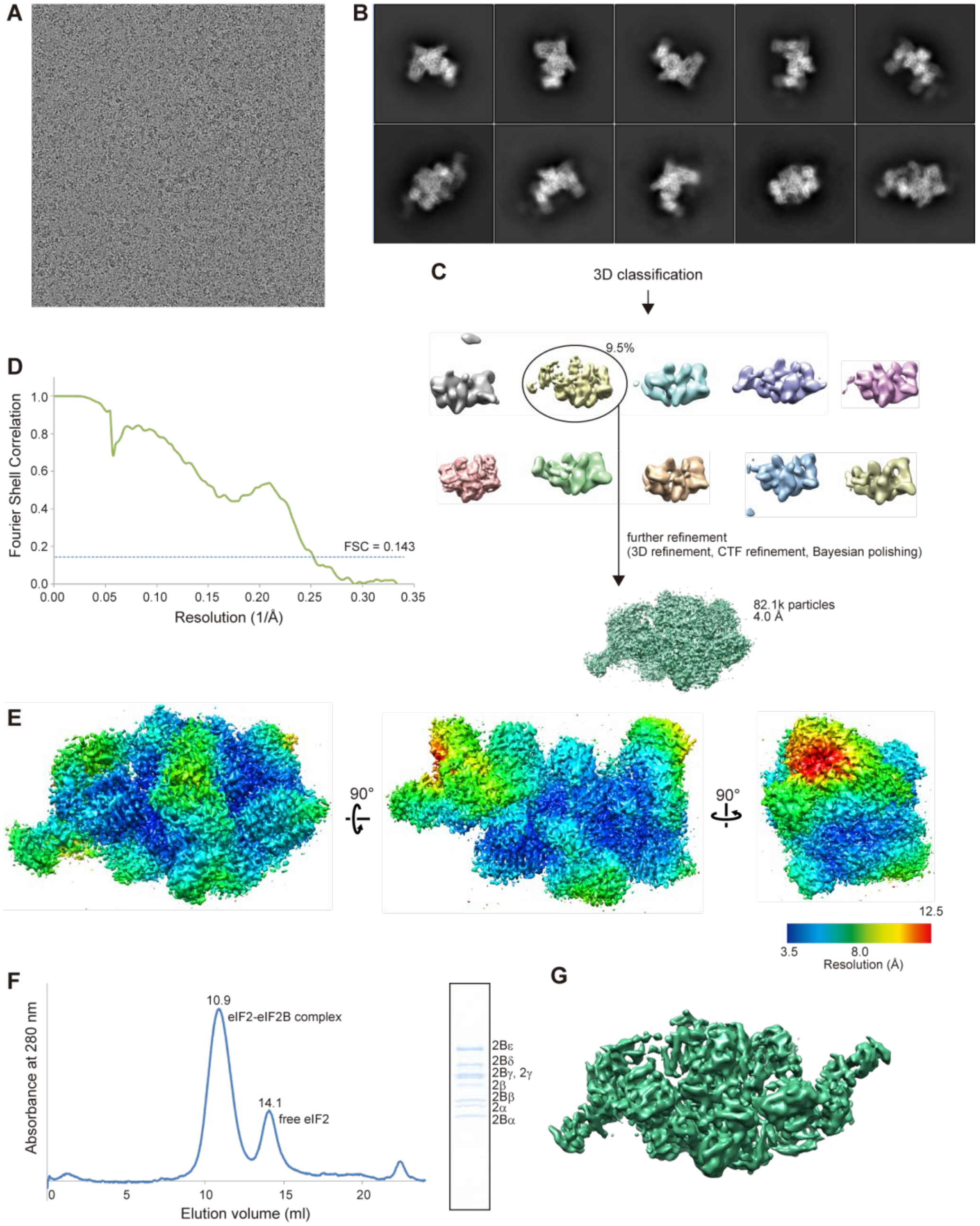

**Fig. S1. Cryo-EM data processing for the eIF2•eIF2B complex.**

(**A**) Representative image of micrographs. (**B**) Representative images of the reference-free 2D class averages of particles. (**C**) Workflow of image processing. (**D**) Fourier shell correlation curve of the cryo-EM map. The curve drops below 0.143 at a resolution of 4.0 Å. (**E**) Local resolution maps from three different views. The density around the G domain of eIF2γ has the lowest resolution. (**F**) Size exclusion chromatography analysis of the mixture of eIF2 and eIF2B with a Superose 6 column. The chromatogram of the absorbance at 280 nm and the Coomassie Brilliant Blue stained SDS-PAGE gel of the peak fraction. (**G**) The 6.8-Å resolution structure of eIF2B in complex with two molecules of eIF2, reconstructed from the sample prepared without detergent.

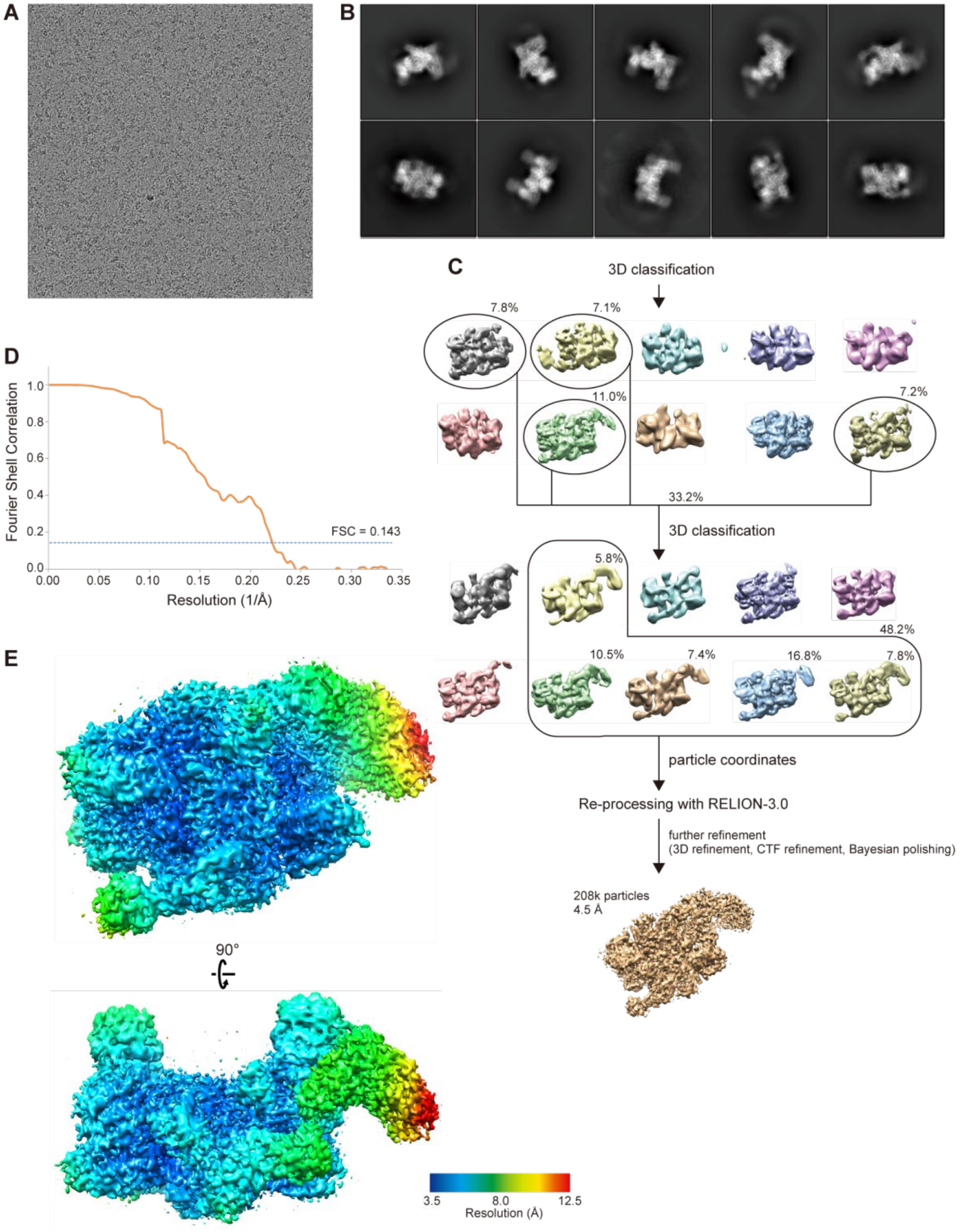

**Fig. S2. Cryo-EM data processing for the eIF2(αP)•eIF2B complex.**

(**A**) Representative image of micrographs. (**B**) Representative images of the reference-free 2D class averages of particles. (**C**) Workflow of image processing. (**D**) Fourier shell correlation curve of the cryo-EM map. The curve drops below 0.143 at a resolution of 4.5 Å. (**E**) Local resolution maps from two different views.

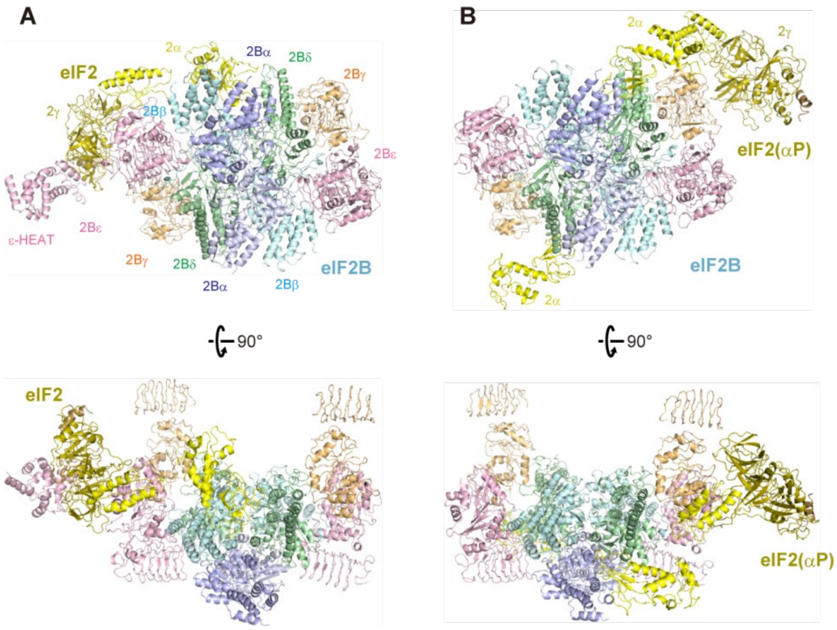

**Fig. S3. Cartoon representations of the human eIF2•eIF2B complexes.**

The cartoon representations of the unphosphorylated eIF2•eIF2B complex (A) and the phosphorylated eIF2(αP)•eIF2B complex (B). The same view and coloring as Fig. 1.

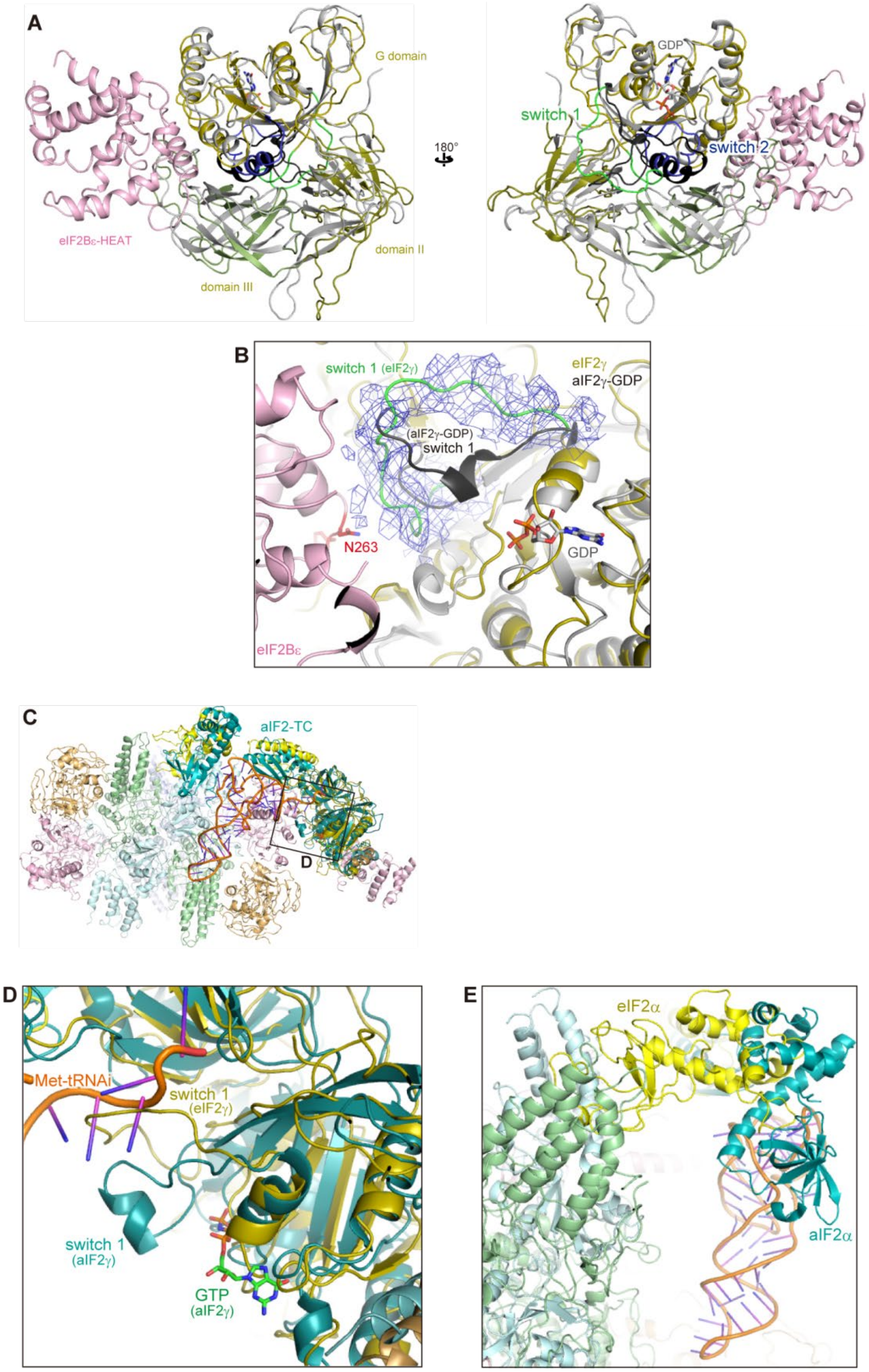

**Fig. S4. The catalytic mechanism of eIF2B.**

(**A** and **B**) The eIF2γ movement induced by eIF2B binding. (A) The movement of three domains of eIF2γ upon eIF2B binding. The G domains of eIF2γ in the human eIF2•eIF2B complex and aIF2γ in the aIF2-GDP complex (grey, PDB ID: 2QMU) (25) are aligned. The switch 1 and 2 regions in eIF2γ are colored green and blue, and those in aIF2γ are colored black, respectively. (B) The EM density map around the switch 1 region of eIF2γ is shown in blue. The same view as Fig. 2B. (**C** to **E**) Structural alignment of eIF2-eIF2B and aIF2-TC. (C) The structures of the eIF2•eIF2B complex and aIF2-TC (PDB ID: 3V11) (28) are aligned with their γ-subunits of a/eIF2. aIF2 is colored teal. (D) Close-up view around GTP and the CCA tail of Met-tRNA$_i$ in the aIF2-TC structure. (E) Close-up view of the a/eIF2α-NTD.

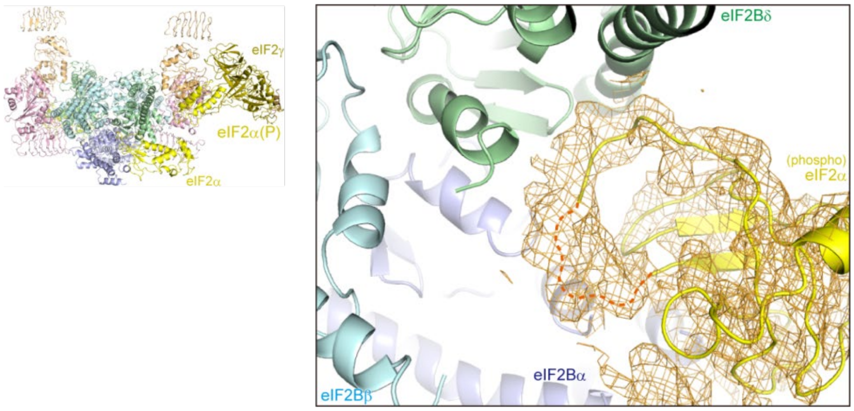

**Fig. S5. EM density map around the Ser51-flanking loop of P-eIF2α in the eIF2(αP)•eIF2B complex.**

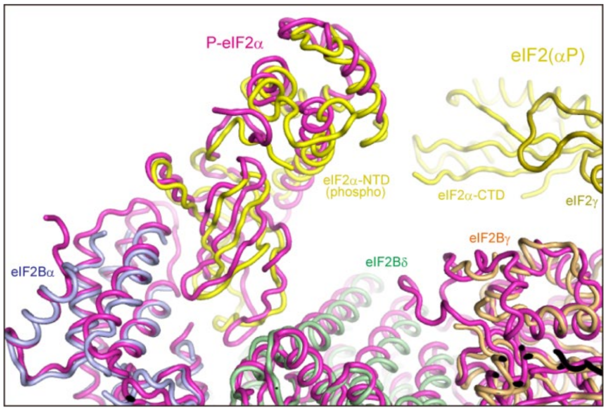

**Fig. S6. Comparison of the P-eIF2α-NTD position in the eIF2(αP)•eIF2B complex and the P-eIF2α•eIF2B complex.**

The structures of the eIF2(αP)•eIF2B complex and the P-eIF2α•eIF2B complex are aligned with their α-subunits of eIF2B. eIF2(αP)•eIF2B is colored as Fig. 1 and P-eIF2α•eIF2B is colored magenta.

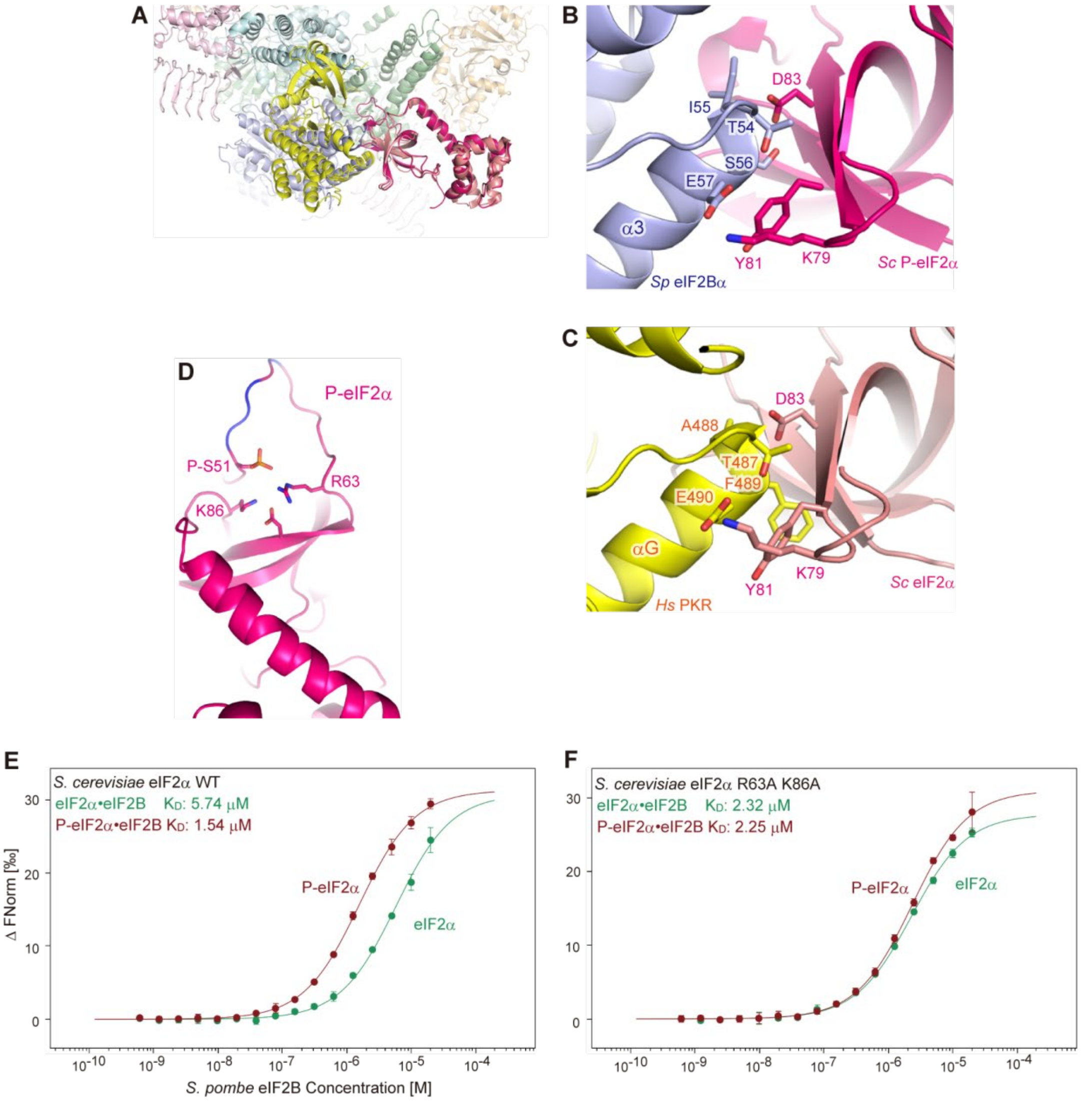

**Fig. S7. The mechanism of eIF2α recognition.**

(**A** to **C**) Comparison of the binding modes of eIF2α in the P-eIF2α•eIF2B complex and the eIF2α•PKR complex. (A) The structures of the P-eIF2α•eIF2B complex and the eIF2α•PKR complex (PDB ID: 2A1A) (30) are aligned with their α-subunits of eIF2. eIF2α and PKR in the eIF2α•PKR complex are colored salmon and yellow, respectively. (B and C) Close-up views of the recognition interface for eIF2α in the P-eIF2α•eIF2B complex (B) and in the eIF2α•PKR complex (C). Key residues for the interactions are shown in stick models. (**D** to **F**) The microscale thermophoresis analysis of the dissociation constants between eIF2α and eIF2B. (D) The structure of P-eIF2α from the P-eIF2α•eIF2B complex. The positions of the phosphorylated residue (P-Ser51) and the substituted residues in this analysis (Arg63 and Lys86) are shown in stick models. (E and F) Dose response curves for unphosphorylated *S. cerevisiae* eIF2α (green) and P-eIF2α (dark red) binding to *S. pombe* eIF2B. The curves for wild type eIF2α (E) and R63A, K86A-substituted eIF2α (F). The dissociation constants ($K_D$) were calculated from triplicate analyses. Error bars represent standard deviations of each data point.

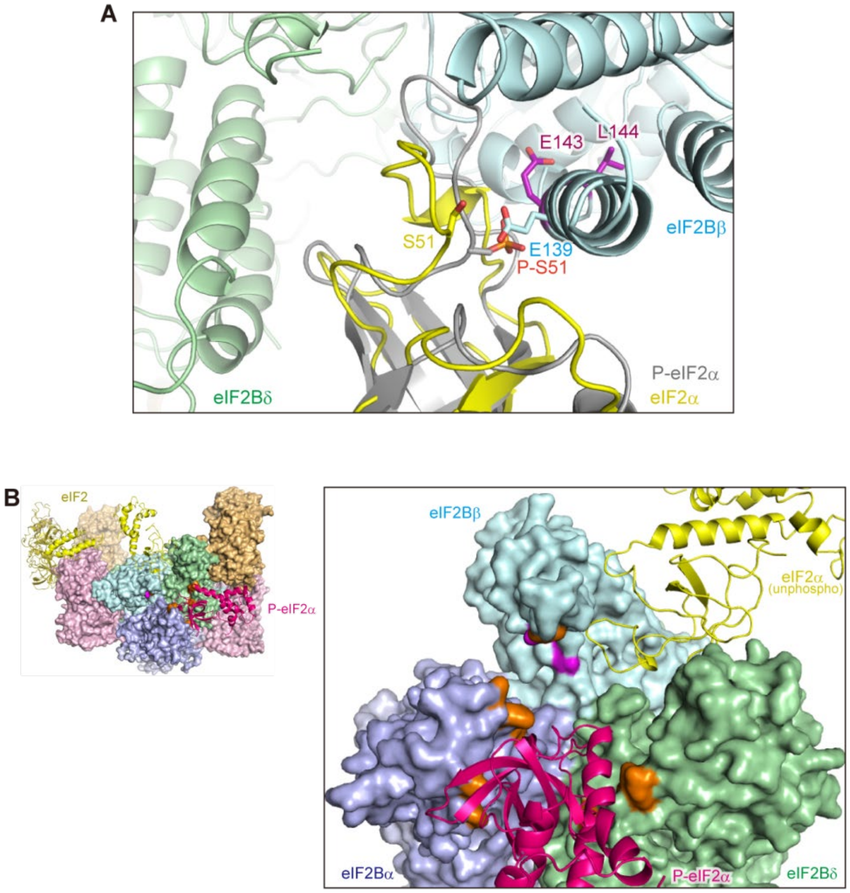

**Fig. S8. Phosphorylation of eIF2α and interaction with eIF2B.**

(**A**) P-eIF2α from P-eIF2α•eIF2B (grey) is overlaid onto eIF2α in the eIF2•eIF2B complex. The same view as Fig. 4A. P-Ser51 of P-eIF2α is shown in a stick model. (**B**) The residues of eIF2B cross-linked with eIF2α in the previous study (14) are colored on the surface model of eIF2B from the P-eIF2α•eIF2B structure. The residues cross-linked with both eIF2α and P-eIF2α are colored orange, and the residues whose cross-links were diminished by the phosphorylation of eIF2α are colored violet. The unphosphorylated eIF2α is modelled by the structural alignment of the eIF2•eIF2B complex and the P-eIF2α•eIF2B complex with their β- and δ-subunits of eIF2B.

| | eIF2•eIF2B | eIF2(αP)•eIF2B |
|---|---|---|
| **Data collection and processing** | | |
| Magnification | 23500 | 23500 |
| Voltage (kV) | 200 | 200 |
| Electron exposure (e–/Å$^2$) | 50 | 50 |
| Defocus range (μm) | -1.5 to -3.1 | -1.5 to -3.1 |
| Pixel size (Å) | 1.47 | 1.47 |
| Symmetry imposed | C1 | C1 |
| Initial particle images (no.) | 825568 | 1482123 |
| Final particle images (no.) | 82123 | 208245 |
| Map resolution (Å) | 3.99 | 4.54 |
| FSC threshold | 0.143 | 0.143 |
| **Refinement** | | |
| Initial model used (PDB code) | 5B04 | 5B04 |
| Model resolution (Å) | 4.10 | 4.59 |
| FSC threshold | 0.5 | 0.5 |
| Map sharpening *B* factor (Å$^2$) | -109.4 | -184.7 |
| Model composition | | |
| Non-hydrogen atoms | 32587 | 32770 |
| Protein residues | 4203 | 4232 |
| Ligands | 0 | 0 |
| *B* factors (Å$^2$) | | |
| Protein | 66.7 | 65.8 |
| R.m.s. deviations | | |
| Bond lengths (Å) | 0.007 | 0.008 |
| Bond angles (°) | 1.189 | 1.239 |
| Validation | | |
| MolProbity score | 2.14 | 2.21 |
| Clashscore | 9.34 | 11.01 |
| Poor rotamers (%) | 0.72 | 0.97 |
| Ramachandran plot | | |
| Favored (%) | 86.0 | 85.4 |
| Allowed (%) | 14.0 | 14.6 |
| Disallowed (%) | 0.0 | 0.1 |

**Table S1. Cryo-EM data collection and image processing.**

| | P-eIF2α•eIF2B | eIF2α•eIF2B |
|---|---|---|
| **Data collection** | | |
| Space group | $P2_1$ | $P2_1$ |
| Cell dimensions | | |
| *a*, *b*, *c* (Å) | 155.58, 207.71, 155.74 | 155.54, 208.13, 155.61 |
| β (°) | 96.96 | 97.14 |
| Resolution (Å) | 50–3.35 (3.55–3.35)* | 50–3.50 (3.71–3.50) |
| $R_{sym}$ or $R_{merge}$ | 0.099 (1.092) | 0.082 (1.278) |
| $I / \sigma I$ | 12.06 (1.25) | 13.03 (1.25) |
| Completeness (%) | 99.7 (98.8) | 99.6 (98.6) |
| Redundancy | 3.49 (3.45) | 3.49 (3.52) |
| $CC_{1/2}$ | 0.998 (0.501) | 0.999 (0.515) |
| **Refinement** | | |
| Resolution (Å) | 49.31–3.35 | 49.31–3.50 |
| No. reflections | 272064 | 238755 |
| $R_{work}$ / $R_{free}$ | 0.221 / 0.253 | 0.224 / 0.260 |
| No. atoms | | |
| Protein | 31766 | 31771 |
| Ligand/ion | 60 | 40 |
| Water | 0 | 0 |
| *B*-factors | 111.5 | 144.1 |
| Protein | 111.5 | 144.1 |
| Ligand/ion | 126.9 | 152.5 |
| R.m.s. deviations | | |
| Bond lengths (Å) | 0.006 | 0.004 |
| Bond angles (°) | 0.633 | 0.587 |

Both datasets were collected from single crystals.

*Values in parentheses are for highest-resolution shell.

**Table S2. X-ray data collection and refinement statistics.**
